# High quality frozen extracts of *Xenopus laevis* eggs reveal size-dependent control of metaphase spindle micromechanics

**DOI:** 10.1101/112334

**Authors:** Jun Takagi, Yuta Shimamoto

**Affiliations:** Quantitative Mechanobiology Laboratory, Center for Frontier Research, National Institute of Genetics; Department of Genetics, School of Life Science, SOKENDAI; AMED-PRIME, Yata 1111, Mishima, Shizuoka 411-8540, Japan

## Abstract

Cell-free extracts from unfertilized *Xenopus laevis* eggs offer the opportunity for a variety of biochemical assays for analyzing essential cell cycle events, such as metaphase spindle assembly. However, extracts’ utility is often hampered by their short storage-stability duration and high quality variation. Here, we report a simple two-step method for preparing frozen egg extracts that retain spindle assembly activity levels that are close to those of freshly prepared extracts. Extract degradation associated with the freeze-thaw process can be substantially reduced by using centrifugal filter-based dehydration and slow sample cooling. Large amounts of frozen extract stocks from single batch preparations allowed us to collect a large number of data in micromanipulation experiments that are intrinsically low-throughput and, hence, clarify correlations between metaphase spindle size and stiffness. We anticipate that our method provides an assay platform with minimized biological heterogeneity and makes egg extracts more accessible to researchers as distributable material.

**Summary:** The authors describe a method for preparing frozen extracts of *Xenopus laevis* eggs that retain spindle assembly activity levels that are close to those of freshly prepared extracts. This allowed for clarifying the correlation between spindle size and stiffness.

## Introduction

Proper assembly of the microtubule-based bipolar spindle is crucial for successful cell division in eukaryotes, and understanding the molecular mechanisms regulating this process has been a major focus in cell biology. Cell-free extracts prepared from the unfertilized eggs of *Xenopus laevis* is a model experimental system that has substantially contributed towards understanding these mechanisms. This system is open to a variety of biochemical manipulations such as fractionation and immunodepletion to identify important proteins required for proper functioning of the spindle (Murray, 1991; Desai et al., 1999; Hannak and Heald, 2006). The assembled spindles can also be targeted for micromanipulation studies, using cantilevers or glass microneedles, thus allowing for an examination of the spindle's structural stability against mechanical perturbations (Itabashi et al., 2009; Gatlin et al., 2010; Shimamoto et al., 2011; Takagi et al., 2013; Takagi et al., 2014). Despite such experimental versatility, this system suffers from several potential drawbacks. The endogenous activity of the extracts is retained for only a short time (typically 2-8 hours from when the eggs are crushed). Moreover, extracts exhibit great variability in quality, mostly due to the eggs’ quality and preparation processes, which often arises as considerable variations in spindle phenotypes, such as variations in size, shape, and response to perturbations (Hannak and Heald, 2006; Grenfell et al., 2016). These discrepancies in quality make it difficult to collect a large number of data sets under a controlled biological setting, particularly for measurements that are intrinsically low-throughput, i.e., that span over a long period and are restricted to examine one sample at a time. Equipment costs, such as the maintenance of frog colonies and purchase of specialized instruments, can also hamper the accessibility of such extracts.

One way to overcome these problems is to prepare frozen extract stocks, the quality of which is even and well characterized. Frozen extracts that are routinely used in laboratories are capable of reconstituting essential cell cycle events such as DNA replication and chromosome organization, but not bipolar spindle assembly (Hannak and Heald, 2006; Gillespie et al., 2012). Classically, the cryopreservation of biological materials such as cells and tissues has been extensively studied with the consensus that the key to successful preparation is the prevention of ice crystal formation in the cytoplasm, which may cause significant damage to organelles and other cellular structures (Fowler and Toner, 2005). A widely used technique for preventing ice crystal formation is to supplement the freezing medium with cryoprotectants such as DMSO. However, these reagents are often toxic at their effective concentrations and difficult to remove from the cytoplasm during sample recovery. Another important technique is to freeze a sample at a controlled speed. Extremely fast cooling (e.g., snap freezing) can be ideal for preventing ice crystal formation; however, this can only be achieved within micron depth from the contact surface. Alternatively, samples can be cooled at a slow speed (e.g., −1°C/min). This method allows the liquid outside the cell to crystalize first, making the cells hypertonic; water then flows out of the cytoplasm through the cell membrane, reducing the probability of ice formation within the cell. However, too much dehydration causes loss of cellular activity because it alters the balance of the cell's chemical composition (Fowler and Toner, 2005). Additionally, samples must be rehydrated for their recovery during thawing. Although such methods have been developed for cells and tissues (Kaiser, 2002; Fowler and Toner, 2005), applying these strategies to egg extracts has not been effective because of their cell-free nature.

Here, we report a simple two-step method for preparing frozen *Xenopus* egg extracts for which the biochemical activity is retained at levels that are close to those of freshly prepared extracts. Using a centrifugal filter membrane and a slow cooling device, an optimal freezing condition can be achieved without a loss of spindle assembly activity. This allowed us to repeat a substantial number of force measurements in bipolar metaphase spindles, which are intrinsically low-throughput and typically allow only a few samples to be examined in each fresh preparation. Through our measurements, we were able to reveal the existence of a scaling mechanism between spindle size and stiffness.

## Results

Cryopreservation studies suggest that a critical problem associated with sample freezing is the formation of ice crystals, and this can be circumvented by dehydration of the cytoplasm, which typically occurs through the plasma membrane when cells are cooled at a slow rate. Inspired by this, we separated the water content in freshly prepared extracts, using a centrifugal filter membrane device (upper panel, Fig. 1A). Briefly, metaphase-arrested fresh extracts were first prepared according to the standard protocol (Desai, 1999), and then loaded into a filter device. A centrifugation for 10 min, using a 100-kDa mesh, resulted in the concentrate and flow-through fractions, which were each transferred to test tubes and separately cooled to −80°C at approximately − 1°C/min. Prior to experimental assays, the frozen concentrate and flow-through fractions were thawed on ice and then combined (lower panel, Fig. 1A). Upon centrifugation, the endogenous protein concentration of the extract, which was initially 80 ± 5 mg/ml as determined by the Bradford method (mean ± S.D., n = 3 preparations), was concentrated to 97 ± 10 mg/ml (Fig. S1A). In contrast, the flow-through fraction was ~1/4 the volume of the loaded extracts and contained only a trace amount of protein (1.8 ± 0.2 mg/ml, ~0.5 % of the total protein content). Upon recovery, the extracts had a protein concentration of 79 ± 5 mg/ml (n = 3), which is comparable to that of the original fresh extracts. An SDS-PAGE analysis showed no prominent loss of certain proteins after the entire procedure (Fig. S1B). We refer to the extracts prepared by this method as filtered frozen extracts.

**Figure 1.**
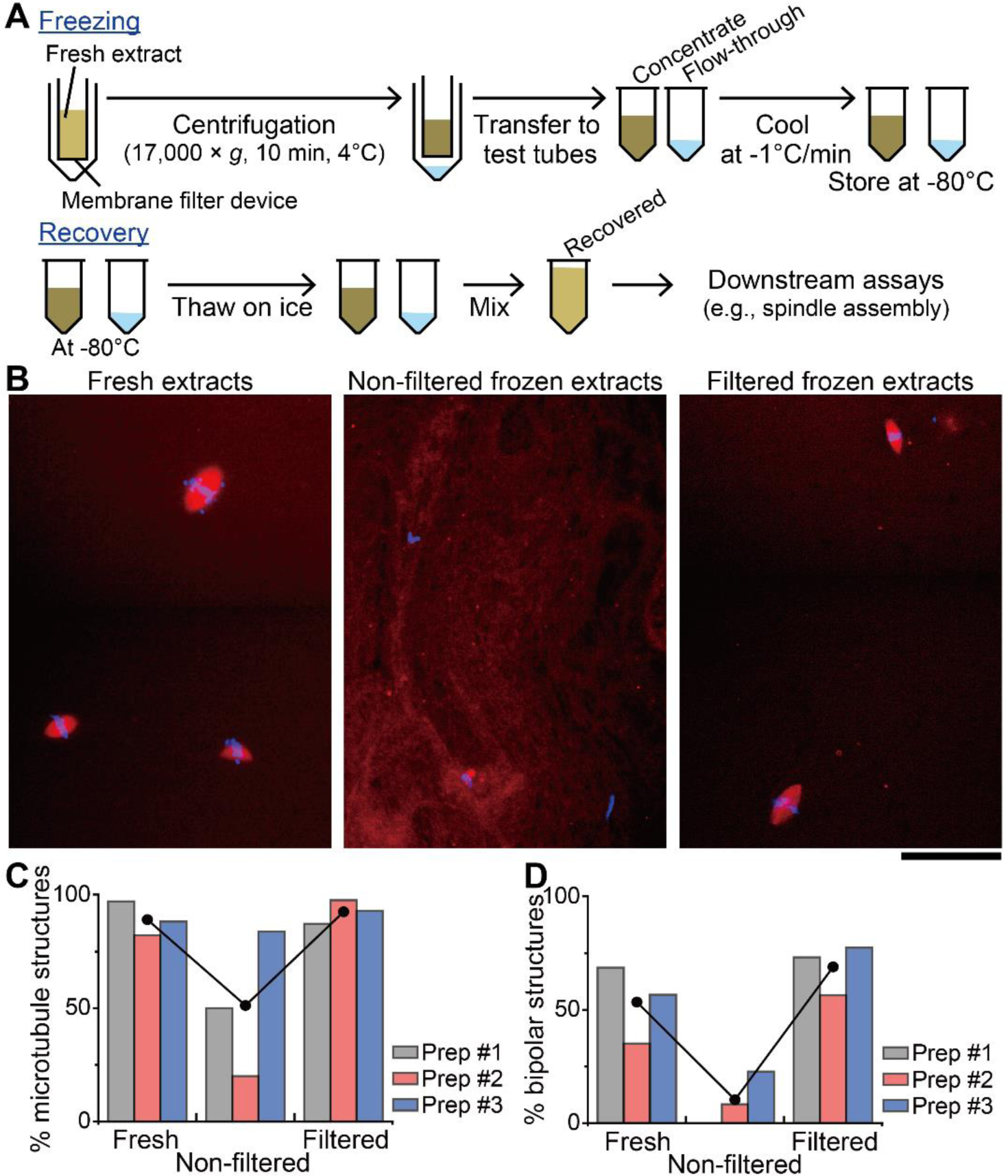
Preparation and use of frozen *Xenopus laevis* egg extracts that retain spindle assembly activity. (**A**) Procedure for preparing frozen *Xenopus* egg extracts that are capable of bipolar spindle assembly. For freezing, freshly prepared extracts (yellow) are loaded into a centrifugal filter device, and then spun at indicated parameters. The resultant concentrate (dark brown) and flow-though (light blue) fractions are each transferred to test tubes and cooled to −80°C at approximately − 1°C/min. For recovery, the frozen fractions are thawed on ice and then combined. (**B**) Representative fluorescence images of microtubule-based structures (red: tetramethylrhodamine-tubulin, 500 nM) assembled around metaphase sperm nuclei (blue: DAPI, 1 μg/ml). Freshly prepared extracts, frozen extracts prepared without filtration, and those prepared with filtration were fixed in squashes after cycling once through interphase and back into metaphase. Scale bar, 100 μm. (**C**) Fraction of sperm nuclei that associated with significant microtubule assembly. Data from three independent preparations are shown (colored bars). Integrated fluorescence intensity of labeled tubulins was analyzed around each sperm nucleus in fresh extracts (n = 69, 90, and 68), non-filtered frozen extracts (n = 64, 60, and 105), and filtered frozen extracts (n = 47, 87, and 43); those that exceeded a certain threshold were scored. (**D**) Fraction of microtubule-based structures that were of bipolar shape, with focused poles. The structures scored in (**C**) were analyzed to examine their bipolarity (n = 67, 74, and 60 for fresh extracts; n = 32, 12, and 88 for non-filtered frozen extracts; n = 41, 85, and 40 for filtered frozen extracts).

To test how much of their endogenous activity the filtered frozen extracts were able to retain, we ran a cycled spindle assembly reaction (Desai, 1999) and examined microtubule-based structures assembled around metaphase sperm nuclei. The structure's phenotypes were analyzed in fixed squashes and compared with those of freshly prepared extracts and frozen extracts prepared without filtration; comparisons were made between extracts that originated from identical preparation batches. We found that the major fraction of sperm nuclei in filtered frozen extracts had prominent microtubule assembly activity that was close to the activity in fresh extracts, and many of the structures were bipolar (left and right panels, Fig. 1B). In contrast, in non-filtered frozen extracts, little or none of those structures were observed (center panel, Fig. 1B). We quantitatively analyzed the extracts’ microtubule assembly activity by counting the number of sperm nuclei that had more than a threshold level of tubulin fluorescence. The analysis was repeated with three independent preparations. We found that in filtered frozen extracts, the fraction of sperm nuclei with significant tubulin signal was as high as that in the fresh extracts in all three preparations (93 ± 5 % versus 89 ± 7 %, mean ± S.D.); whereas, in non-filtered frozen extracts, two of three preparations showed considerable reduction in that activity (<50 %) (Fig. 1C). In filtered frozen extracts, 69 ± 11 % of these structures were of the typical bipolar shape with focused poles, which was comparable to or even slightly higher than that in the fresh extracts (53 ± 17 %) (Fig. 1D) and in those reported previously (50-80 %; Budde et al., 2001; Miyamoto et al., 2004; Helmke and Heald, 2014). In contrast, in non-filtered frozen extracts, only ~10 % of the assembled structures were bipolar (10 ± 11 %) (Fig. 1D). In all experimental conditions, the remaining fraction predominantly comprised monopolar, multipolar, or disorganized structures (Fig. S2A). The bipolar spindles in filtered frozen extracts maintained their overall morphology and equatorial chromosome alignment for >20 min, indicating that the extracts retained cell cycle arrest at metaphase (Fig. S2, B and C). Therefore, our filtration method significantly improves the frozen extracts’ capability to assemble the bipolar metaphase spindle.

**Figure 2.**
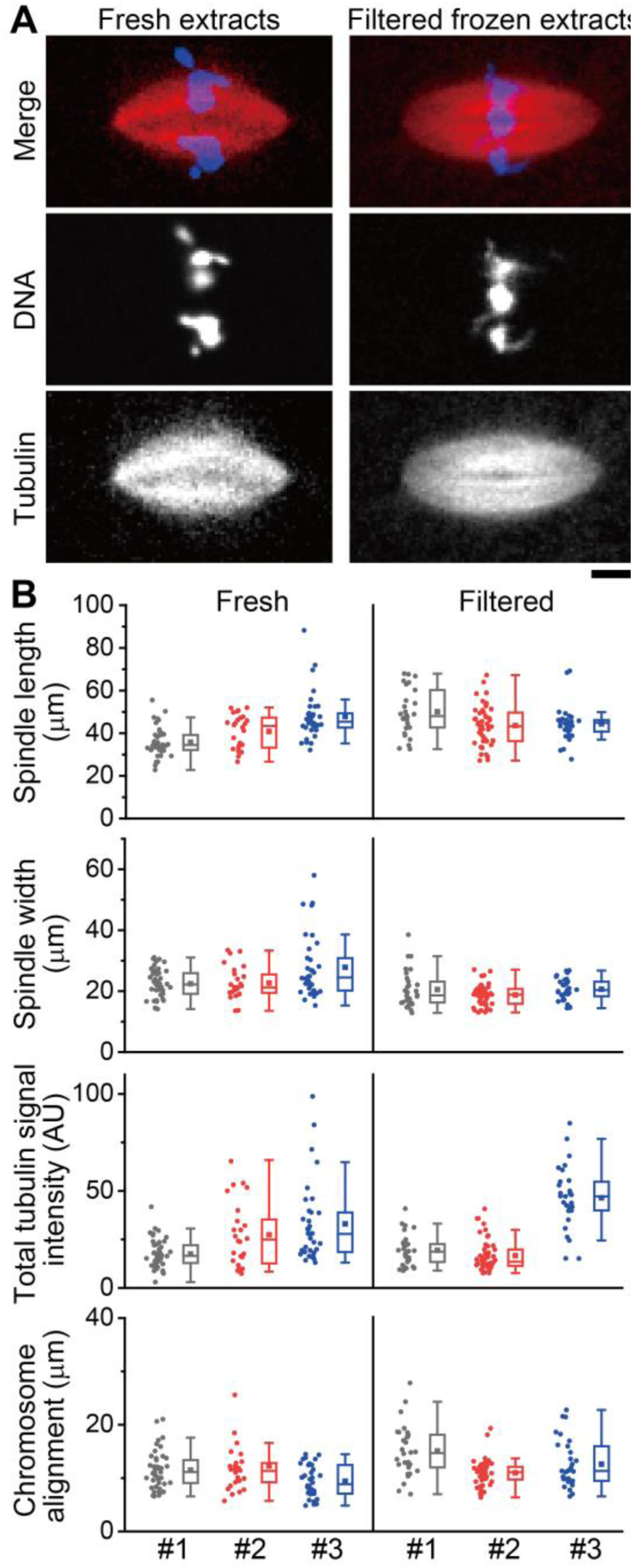
Metaphase spindle morphology in fresh and filtered frozen extracts. (**A**) Representative fluorescence images of bipolar metaphase spindles (red: tetramethylrhodamine-tubulin, blue: DAPI) assembled in fresh extracts or filtered frozen extracts. Scale bar, 10 μm. (**B**) Distributions of spindle morphology parameters. Bipolar structures scored in Fig. 1D were analyzed to determine individual spindle length, width, total tubulin signal intensity, and chromosome alignment for fresh (left columns; n = 46, 26, and 34 spindles) and filtered frozen extracts (right columns; n = 30, 48, and 31 spindles). Data from three independent preparations are shown in colored box charts with individual plots. Within the box charts, the solid squares indicate mean values, and the horizontal lines indicate median values. The bottoms and tops of the boxes indicate first and third quartiles, respectively. Whiskers show highest and lowest values within 1.5 times the interquartile range. For statistical analysis results, see Table S1.

To quantitatively assess the morphology of the assembled structures, we analyzed the length, width, tubulin quantity, and chromosome alignment of bipolar spindles, using an automated image analysis pipeline (Grenfell et al., 2016) (Fig. 2A). In agreement with previous study (Grenfell et al., 2016), the morphology of individual spindles in fresh extracts had broad distributions within each preparation (left three columns, Fig. 2B), and their median values largely varied between preparations (horizontal lines in the box charts). For example, the median spindle lengths analyzed in three independent preparations were 34.5, 43.3, and 45.4 μm (n = 46, 26, and 36, respectively). For quantities of other parameters, see Supplementary Table S1. The morphology parameters analyzed for filtered frozen extract spindles also had large variations in their distributions and median values (right three columns, Fig. 2B). For example, the median spindle lengths in three independent preparations were 48.0, 43.2, and 44.9 μm (n = 30, 48, and 31, respectively). Notably, although one of the three preparations showed relatively large variance in median spindle length between filtered frozen extracts and their original fresh extracts (39 %, Prep #1 in Fig. 2B), the remaining two preparations yielded significantly small deviations in their median spindle lengths (0.2 % and 1.1 %, Prep #2 and #3 in Fig. 2B, respectively). A typical variation of median spindle length that arises upon each fresh preparation is ~10-20 % (14 % for n = 3 independent preparations in this study; 11, 13, and 17 % for n = 26, 19, and 5 preparations, respectively, in studies conducted by different experimenters in Grenfell et al., 2016). The analysis was also performed for other spindle morphology parameters, and the differences between filtered frozen extracts and their original fresh extracts were at most ~2-fold of the typical variations seen among fresh preparations (Supplementary Table S2). Therefore, variations in spindle morphology between filtered frozen extracts and their original fresh extracts are, if any, comparable to those that arise during fresh extract preparation.

Successful cryopreservation requires an empirical determination of optimal dehydration, cooling, and recovery (Fowler and Toner, 2005). We therefore examined factors that might affect the quality of frozen extract stocks. First, we examined the speed of sample freezing and found that the extracts should be cooled at a slow speed (e.g., −1°C/min). Snap-freezing the concentrate fraction resulted in significant loss of spindle assembly activity (Fig. S3, A and B). Second, we examined the speed of sample thawing and found that spindle assembly activity was not significantly affected by whether the frozen extracts were thawed on ice or at 16 °C (Fig. S3, C and D). Third, we tested the requirement of extract re-hydration and confirmed that the entire flow-through fraction has to be added back to the concentrate to restore the extract's endogenous activity. Without this rehydration, massive microtubule polymerization occurred all throughout the cytoplasm (Fig. S3, E and F). Fourth, we examined the period of centrifugation and found that it should be limited to less than 10 min. Increasing the centrifugation time resulted in larger flow-through volume and thus more dehydration of the extracts (Fig. 3A). The spindle assembly activity was improved accordingly, but excess centrifugation impaired extracts’ quality (Fig. 3, B and C). Finally, we compared different filter mesh sizes and found that a 100-kDa mesh should be optimal. Preparation with smaller mesh sizes (e.g., 10 kDa) resulted in less amount of protein elution in the flow-through (<0.2 mg/ml, versus 1.8 mg/ml with a 100-kDa mesh). The concentration rate of extracts was consequently reduced by ~3 % (Fig. 3E). Although the microtubule assembly activity was nearly independent of the filter mesh size (Fig. 3F), extracts prepared using a 100-kDa filter constantly yielded higher fractions of bipolar spindle phenotype (Fig. 3G). We note that in contrast to the marked sensitivity of spindle assembly probability to the preparation parameters, the bipolar spindle morphology was quite robust to those parameters (Fig. 3, D and H). Together, these results highlight the importance of dehydration level and freezing speed in our preparation procedure to assemble bipolar spindles at numbers similar to those that can be achieved in fresh extracts.

**Figure 3.**
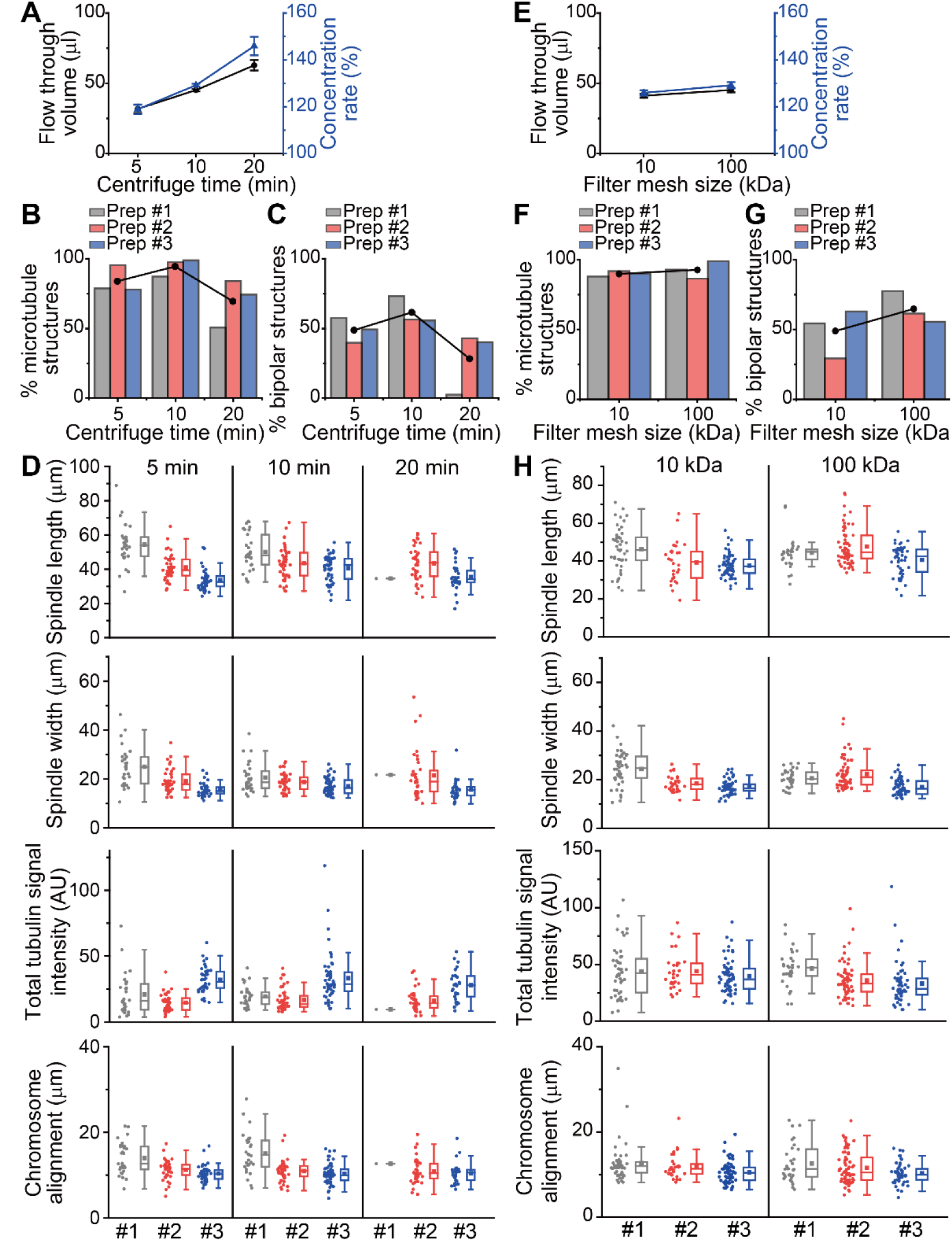
Optimization of the frozen preparation procedure. Dependencies of the extract's spindle assembly capacity on centrifugation time (**A-D**) and filter mesh size (**E-H**). Data from three independent preparations for each condition are shown (colored bars and plots). (**A, E**) Concentration rate (blue triangles, in % of the original extract volume) estimated from the volume of flow-through (black circles, in μl) obtained with indicated centrifugation time (**A**) or filter mesh size (**E**). Plots are presented as mean ± S.D. (n = 3). (**B, F**) Fraction of sperm nuclei that were associated with significant microtubule assembly. Analysis was performed as in Fig. 1C. The structures analyzed were n = 66, 108, and 109 for 5-min spin; n = 47, 87, and 98 for 10-min spin; and n = 77, 100, and 101 for 20-min spin, respectively, in (B), and n = 100, 111, and 111 for 10-kDa filter and n = 43, 126, and 98 for 100-kDa filter, respectively, in (**F**). (**C, G**) Fraction of microtubule-based structures that were of bipolar shape with focused poles. Analysis was performed as in Fig. 1D. The structures analyzed were n = 52, 103, and 85 for 5-min spin; n = 41,85, and 97 for 10-min spin; and n = 39, 84, and 75 for 20-min spin, respectively, in (**C**), and n = 88, 102, and 100 for 10-kDa filter and n = 40, 109, and 97 for 100-kDa filter, respectively, in (**G**). Black circles in (**B**), (**C**), (**F**), and (**G**) are mean values in each preparation condition. (**D, H**) Distributions of spindle morphology parameters. Bipolar structures scored in (**C**) (n = 30, 41, and 42 for 5-min spin; n = 30, 48, and 54 for 10-min spin; and n = 1,36, and 30 for 20-min spin, respectively) and (**G**) (n = 48, 30, and 63 for 10-kDa filter and n = 31,67, and 54 for 100-kDa filter, respectively) were analyzed to determine individual spindle length, width, total tubulin signal intensity, and chromosome alignment in each preparation condition. Box chart representations are as in Fig. 2B.

Upon establishing the method for preparing frozen extract stocks with retained spindle assembly activity, we examined the mechanical response of metaphase spindles to stretching force applied and its dependency on overall spindle size. To this end, metaphase spindles were assembled in filtered frozen extracts and stretched by the microneedle-based setup we developed previously (Shimamoto et al., 2011; Takagi et al., 2014). Each measurement process typically takes >10 min, and 5-10 spindles can be examined in a single spindle assembly reaction. To collect a large number of data (n > 50), the spindle assembly reaction was performed using several frozen stocks originating from a single batch preparation (Fig. 4A), and results were confirmed across several preparation batches. Fig. 4B shows typical time-lapse images acquired during the measurement. A single metaphase spindle was captured using a pair of glass microneedles, the tips of which were inserted into each half of the spindle (0 s, Fig. 4B). Application of an outward stretching force, which was performed by moving the tip of one microneedle (blue arrowheads, Fig. 4B) away from the other, resulted in elongation of the spindle (4 s, Fig. 4B). A tension developed across the bipolar structure, which was revealed by the displacement of the force-calibrated flexible microneedle tip from the equilibrium position (red arrowheads and the broken line, Fig. 4B), and increased as the spindle was stretched (8 s, Fig. 4B). The spindle pole was gradually disorganized as the amount of stretching force increased, and then eventually split apart (white arrows, 12-16 s, Fig. 4B). As a result, the microneedle tip was detached from the spindle and returned to the equilibrium position (16 s, Fig. 4B). A quantitative analysis of force and spindle deformation revealed that the tension first increased in an approximately linear fashion with increasing spindle length and decreasing spindle width (2-9 s, Fig. 4C), and then reached near plateau before the pole was split (~11 s, Fig. 4C). Based on the time recordings we could generate a force-extension plot (Fig. 4D), from which two mechanical properties of the spindle were determined: 1) the stiffness of the entire spindle structure, as analyzed based on a linear regression within the initial 5 % range of extension (red line, Fig. 4D), and 2) the mechanical strength of the spindle pole against motion of a microneedle along the spindle's long axis, as analyzed based on the peak force that appeared at near the maximum spindle extension (blue line, Fig. 4D).

**Figure 4.**
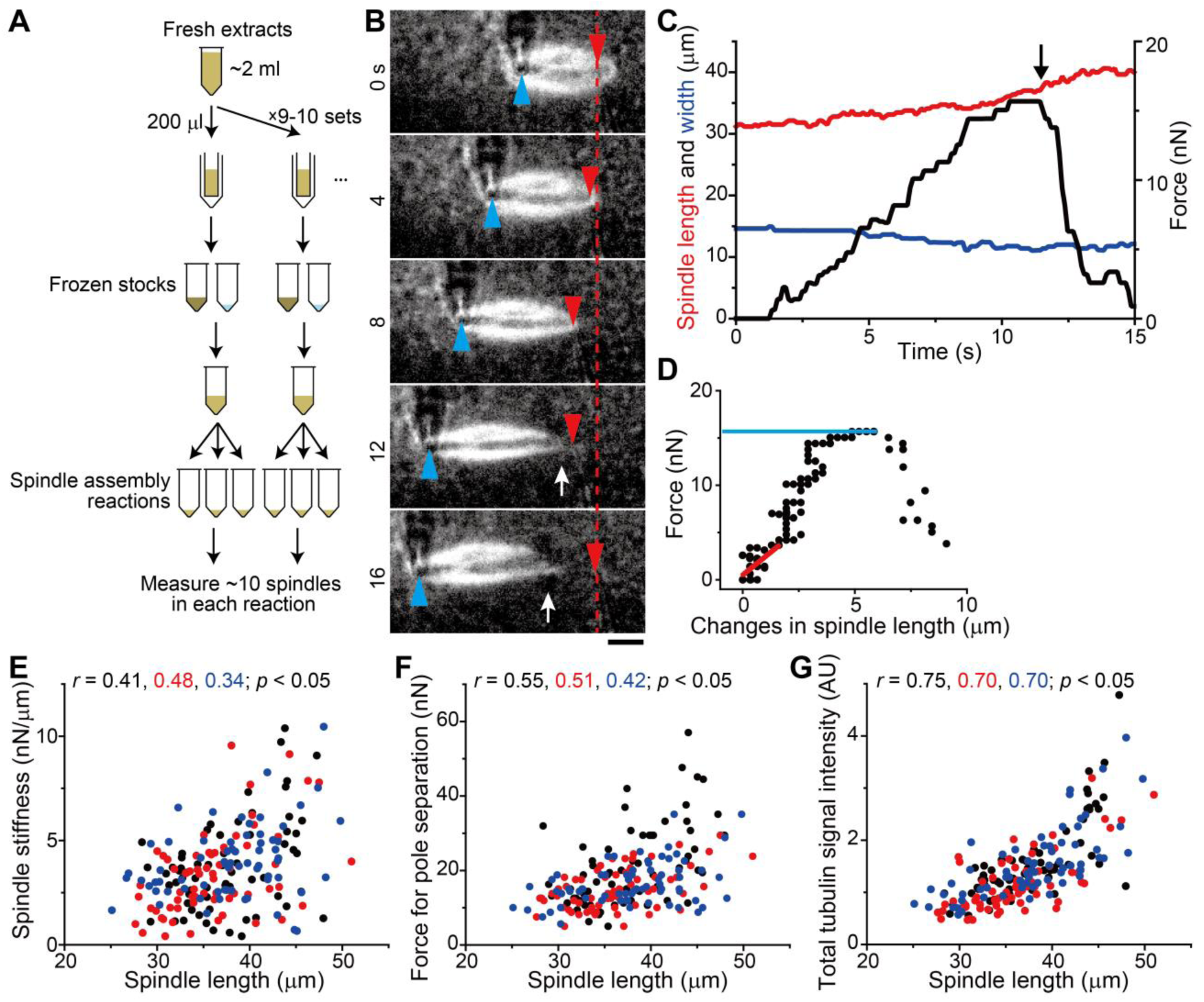
Metaphase spindle micromechanics and their size-dependencies analyzed in filtered frozen extracts. (**A**) Schematic showing the procedure for the batch preparation of filtered frozen extracts used for the assay. Freshly prepared extracts of >2 ml volume are split into aliquots (200 μl each) and subjected to frozen extract preparation, as in Fig. 1A. Each frozen stock is thawed for recovery just prior to use. Data from several days of experiments performed with the same preparation batch are pooled and analyzed. The typical number of data acquired per each reaction is ~10. (**B**) Time-lapse images of a bipolar metaphase spindle assembled in filtered frozen extracts and stretched by a dual microneedle-based setup. Confocal fluorescence (tetramethylrhodamine-labeled tubulin, 500 nM) and bright-field images acquired simultaneously during the time-lapse are overlaid (time stamp: in seconds). In the images, the manipulation microneedle tip (marked in blue) was moved at 2 μm/s. Tension that developed across the spindle was monitored based on the displacement of the force-sensing flexible microneedle tip (marked in red, stiffness: 1.9 nN/μm) from its equilibrium (dashed line). White arrows highlight disorganization and breakage of the spindle pole. Scale bar, 10 μm. (**C**) Changes in spindle length (red), width (blue), and magnitude of force (black) analyzed in (**B**) over time. The black arrow indicates the time at which the spindle pole was split apart. (**D**) Force-extension plots. Changes in spindle length during the time-course in (**C**) are plotted against the magnitude of developed force. Linear regression was performed within 5 % extension range (red solid line) to determine the whole spindle stiffness. Plateau of the force-extension plot (blue dotted line) was used to determine the pole-splitting force. (**E, F**) Dependencies of spindle stiffness (**E**) and pole-splitting force (**F**) on spindle size. A total of n = 65, 66, and 67 spindles were examined in three independent preparations (black, red, and blue circles, respectively) and plotted against each spindle length measured prior to stretch. (**G**) Dependencies of spindle microtubule amount on spindle size. Integrated fluorescence signal intensity of dye-labeled tubulin within each spindle was analyzed for the data set in (**E**) and (**F**). The tubulin images acquired prior to micromanipulation procedure were used for analysis.

We repeated the measurement for a total of n = 63 spindles from a single batch preparation and found that the whole spindle stiffness had a positive correlation with the initial spindle length measured prior to stretch (black circles, Pearson's *r* = 0.41, Fig. 4E).

We repeated this analysis with two additional independent extract preparations and confirmed that the result was consistent among preparations (red and blue circles, Fig. 4E) (Pearson's *r* = 0.48, and 0.34, respectively, *p* < 0.05). Further, the average spindle stiffness was comparable to that measured in fresh extracts in our previous work (4.8 ± 1.7 nN/μm, n = 18 spindles) (Takagi et al., 2014). In addition, we found that the mechanical strength of the spindle pole also had a positive correlation with the spindle length (Fig. 4F; *r* = 0.55, 0.51, and 0.42, *p* < 0.05). Analyzing fluorescence imaging data revealed that the amount of spindle microtubules, as determined based on the signal intensity of dye-labeled tubulins within the bipolar structure, was higher in larger spindles (Fig. 4G; *r* = 0.75, 0.70, and 0.70, *p* < 0.05). Taken together, our assay reveals a scaling mechanism between the metaphase spindle stiffness, size, and amount of microtubules.

## Discussion

We have demonstrated the preparation and use of frozen *Xenopus* egg extracts that possess an endogenous level of spindle assembly activity, and have determined the optimal preparation parameters. Importantly, our method does not rely on cryoprotectants such as DMSO, which enables us to minimize the disturbance in the extracts’ biochemical activity such as microtubule assembly dynamics. Our method thus fulfills the yet-to-be-accomplished process that frozen extract preparation has been lacking, and provides a robust experimental platform for a broad range of applications related to cell division.

Our determination of optimal preparation parameters predicts that the degree of dehydration is an essential part of successful preparation, which agrees with the major cryopreservation technique for cell suspensions. For cells, the freezing speed is a controlled parameter and dehydration occurs in a passive manner depending on cellular characteristics such as permeability of the cell membrane and salt concentration of the cytoplasm. Our method employs centrifugal force to dehydrate extracts and is, therefore, able to control the amount of water removal by altering the spin time, or g-force. Together with analyses of cytoplasmic ultrastructure and composition, our method may help elucidate the physical and quantitative mechanisms of freezing biological materials.

Combined with a microneedle-based force measurement assay and high-resolution fluorescence imaging, our frozen extracts provide important insights into how different-sized metaphase spindles respond to external force. Our assay reveals the existence of a scaling mechanism between the spindle's size and its stiffness. If we assume that the local elasticity of a spindle per unit length is constant and these units are connected in series along the longitudinal axis of the spindle, its total stiffness should be inversely proportional to its length. We previously reported that the volume of *Xenopus* extract spindles exponentially increases with spindle length such that the aspect ratio of the bipolar structure is maintained constant (Takagi et al., 2013). One plausible explanation is that this allometric effect allows for an integrated conjugation of local elastic units in both the long and short axes of the spindle, leading to the size-dependent increase of overall spindle stiffness. The spindle has an ability to dynamically change its size and shape in response to physical and biochemical conditions in the environment (Dumont and Mitchison, 2009; Mitchison et al., 2015; Levy and Heald, 2016; Kapoor, 2017). The scaling mechanism we found here suggests a functional importance of controlling spindle size to ensure the mechanical stability of this essential cytoskeletal structure.

Over the past 30 years, since its development and first use (Lohka and Masui, 1983), the *Xenopus* extract system has been contributing to the discoveries of many essential proteins and pathways for cell cycle events (Hannak and Heald, 2006; Gillespie et al., 2012). Our present method provides a platform for obtaining variable information under minimized sample heterogeneity, enables the standardization of samples, and provides a shippable supply source for researchers. These features will allow this system to be expanded for broader use in the scientific community, and thus contribute to promoting scientific discovery.

## Materials and methods

### Preparation of *Xenopus* egg extracts

Cell-free extracts of unfertilized *Xenopus laevis* eggs arrested at metaphase were prepared as described previously (Desai et al., 1999), except for the use of energy mix of 50× instead of 20×. For preparation of filtered frozen extracts, freshly prepared extracts were loaded into centrifugal filter devices (200 μl per column) (UFC510024 or UFC501024, Millipore) and spun at 17,000 × *g* for 10 min at 4°C in a tabletop centrifuge (5424R, Eppendorf). Following centrifugation, the filter devices were placed upside down in collection tubes and spun at 2,000 × *g* for ~10 s to transfer the concentrate to the tubes. The flow-through fractions were transferred to separate 1.5-ml test tubes using low adhesive pipette tips. These test tubes were placed in a tube cooler (Chillette 12, Denville Scientific Inc.), which was pre-cooled at 4°C, and then frozen at a cooling rate of approximately −1°C/min in a − 80°C freezer. Prior to the spindle assembly reaction, the frozen concentrate and flow-through fractions were thawed on ice and then combined by reintroducing the entire flow-through fraction to the concentrate fraction. The recovered extracts were used for assays following 20-min incubation on ice. Non-filtered frozen extracts, which were used as controls, were prepared in the same manner as above but without centrifugal filtration.

### Preparation of spindle samples

Metaphase spindles were assembled by cycling the extracts with demembranated *Xenopus laevis* sperm nuclei (~400 nuclei/μl) once through interphase and back into metaphase (Desai et al., 1999). The reaction was performed with 20 /μl volume at 16°C in 1.5-ml test tubes. Fixed samples were prepared 90 min after the start of spindle assembly, each with 4 μL of fixation buffer (125 mM Hepes, 2.5 mM EDTA, 2.5 M NaCl, 50 mM KCl, 25 mM MgCl2, 50 mM CaCl2, 10 % formaldehyde, 60 % Glycerol, and 1 /μg/ml DAPI) and 2 /μL of extract reaction supplemented with 500 nM tetramethylrhodamine-tubulin, squashed between a clean slide and a 18×18 mm coverslip. Live spindle samples for micromanipulation experiments were prepared by performing the cycling reaction as above, but with 500 nM tetramethylrhodamine-tubulin and 250 nM SYTOX green.

### Force measurement assays

Mechanical properties of extract spindles were examined using the microneedle-based setup we developed previously (Shimamoto et al., 2011; Takagi et al., 2014). Metaphase extracts (volume: 4 /μl) were placed on a passivated coverslip and covered with mineral oil. The coverslips used were siliconized and pre-coated with Pluronic F-127 (Gatlin et al., 2010). A pair of force-calibrated microneedles, which were prepared according to a previously described method (Shimamoto and Kapoor, 2012) and were each held by three-axis micromanipulators (MHW-3, Narishige), was inserted into each half of a single bipolar spindle. The tip of one microneedle was sufficiently stiff (>1000 nN//μm), to allow for the controlled micromanipulation of spindles, whereas the tip of the other microneedle was much more flexible (stiffness *k* = 1.9 nN//μm), to measure the amount of applied force. The mechanical strengths of the entire spindle structure and its poles were examined by moving the stiff microneedle away from the flexible microneedle along the spindle's pole-to-pole axis at 2 /μm/s. The microneedle motion was controlled using a closed-loop piezo driver (E-665, Physik Instrumente) and a piezo actuator (P-841.20, Physik Instrumente), which was attached to the microneedle's base. The measurements were performed in a temperature-controlled room at 18 ± 1°C and completed within 60 min after spindle assembly, over which no detectable changes in spindle mechanics and morphology were observed.

### Microscopy

Fluorescence imaging of extract samples was performed using an inverted microscope (Ti, Nikon) equipped with a 20× objective lens (Plan Apo, 0.75NA, Nikon), piezo-driven objective scanner (P-725.4CD, PI), sCMOS camera (Neo 4.1, Andor), motorized X-Y stage (MS-2000, Applied Scientific Instrumentation), epi-fluorescence illuminator (Intensilight, Nikon), and spinning-disk confocal unit (CSU-X1, Yokogawa). The epi-illuminator was used with fluorescence filter sets (TxRed-4040C-000 and DAPI-1060B-000, Semrock) for imaging fixed samples. The spinning-disk confocal unit was used with two diode lasers (488 nm and 561 nm, OBIS, Coherent) for live spindle imaging. The entire microscope system was connected to a computer for automated image acquisition using NIS-Elements software (ver. 4.50, Nikon). Fixed samples were subjected to raster-scanning to obtain large images (~10 mm × ~11 mm area per chamber). Live spindle samples were subjected to confocal fluorescence imaging in a streaming mode (exposure time: 100 ms).

### Biochemical characterization

The protein concentration of extracts was determined using the Bradford assay (#500-0006, Bio-Rad) calibrated with the BSA standard. The overall composition of extract proteins was analyzed using SDS-PAGE (4-12 % Tris-glycine, Invitrogen) and Coomassie blue stain (SimplyBlue SafeStain, Invitrogen), according to the manufacturer's protocol. A gel image was acquired using an automated gel imaging instrument (Gel Doc EZ Gel Documentation System, Bio-Rad).

### Data Analysis

Classification of spindle phenotypes, i.e., bipolar, monopolar, multipolar, or disorganized structure, was performed in ImageJ (version 1.49p) by visual inspection of microtubule images acquired around metaphase sperm nuclei in fixed squashes. Quantitative analysis of spindle morphology parameter was performed using a previously published image analysis pipeline (Grenfell et al., 2016). Large aggregates of DNA signals were excluded from analyses.

Spindle length and width in micromanipulation experiments were determined using fluorescence time-lapse images of microtubules, by performing line-scan analyses (line width = 1.6 μm) along the spindle's long axis and short axis, respectively. Typical linescan profile exhibited a bell-like shape with flattened top, from which the edges of the spindle structure was determined at the single pixel resolution by referencing a manually defined threshold that was set at a constant value across the time-lapse images. The magnitude of developed force (*F*) was determined based on the displacement of the force-calibrated microneedle's tip from its equilibrium position (*Δx*), according to *F* = *k · Δx*.

Stiffness of the metaphase spindle was determined based on the relationship between the magnitude of force and the change in spindle length, by performing a linear regression analysis within the initial 5 % increment of spindle length change. The mechanical strength of metaphase spindle poles was determined based on the relationship between the force and the spindle length change, which typically showed a gradual rise followed by a sudden drop, forming a single peak in response to a constant motion of microneedle-based stretch. The critical breakage force was determined at the peak position in the force-extension plot, at which spindle poles showed visible splitting or breakage in fluorescence time-lapse images.

The amount of spindle microtubules in micromanipulation experiments was estimated using fluorescence images of microtubules acquired prior to microneedle insertion. In images acquired for each spindle, a small ROI (Region of Interest; ~50 μm × ~50 μm) and a larger ROI (~100 μm × ~100 μm) were drawn to cover the entire spindle structure. The area and total tubulin signal intensity in the small ROI (*S*_small_ and *I*_small_, respectively) and those in the larger ROI (*S*_large_ and /_large_, respectively) were then obtained. The total tubulin signal intensity in the spindle structure (*I_spindle_*) was determined using these parameter values, according to *I_spindie_ = I*_small_ - *I*_bg_ = *I*_small_ - *S*_small_ · (*I*_large_ − *I*_small_)/(*S*_large_ − *S*_small_). Here, *I* _bg_ is the total background signal intensity in the small ROI, which was estimated based on the signal intensity within the larger ROI but outside the small ROI.

Statistical analyses were performed using a Pearson correlation analysis (Fig. 4, E-G) and an unpaired Student's *t* test (Fig. S1A) in Origin 2016.

## Supplemental materials

Fig. S1 shows the protein concentration and composition of freshly prepared extracts and filtered frozen extracts. Fig. S2 shows non-bipolar phenotypes of microtubule structures and stability of chromosome alignment in filtered frozen extracts. Fig. S3 shows additional data for optimizing the frozen extract preparation. Tables S1 and S2 show statistical data of the spindle morphology parameters in fresh and filtered frozen extracts.

## Acknowledgements

We thank Dr. Masato Kanemaki (National Institute of Genetics) for kindly sharing instruments for method optimization and Dr. Akatsuki Kimura and his laboratory members (National Institute of Genetics) for helpful discussions. This work was supported by JSPS Postdoctoral Fellowship (J.T.), JSPS Grant-in-Aid 16H06166 and 15K14515, and AMED-PRIME (Y.S).

## Author contributions

J.T. and Y.S. designed the experiments and wrote the manuscript; J.T. performed the experiments and analyzed the data.

